# SegmentQTL: Identifying genetic variants influencing molecular phenotypes in copy number–driven cancers

**DOI:** 10.1101/2025.07.28.667150

**Authors:** Samuel Leppiniemi, Daria Afenteva, Yilin Li, Giulia Micoli, Kari Lavikka, Susanna Holmström, Déborah Boyenval, Jaana Oikkonen, Sampsa Hautaniemi, Taru A. Muranen

## Abstract

**Motivation:** Molecular quantitative trait loci (molQTL) analysis links genetic variants to molecular phenotypes, such as gene expression, but existing tools do not account for the structural complexity of copy number–driven cancers. High genomic instability of these cancers leads to chromosomal breaks (breakpoints), which disrupt the physical connection between genes and adjacent regulatory elements. Standard molQTL methods are unable to accommodate breakpoint information and would therefore indiscriminately test associations across breakpoints, leading to spurious signals and reduced biological relevance. To address these challenges, we developed SegmentQTL, a segmentation-aware molQTL analysis tool, designed to improve the accuracy of association testing in unstable cancer genomes by incorporating sample-specific break-point information.

**Results:** SegmentQTL applies an integrated purifying filtering step that removes associations spanning breakpoints, ensuring that only variants within the same segment as the phenotype are tested. This prevents false discoveries and reduces background noise. We evaluated SegmentQTL on selected genes from stable and unstable genomic regions and compared its results with a previously published state-of-the-art tool. In stable regions, SegmentQTL produced similar results, validating its approach. In unstable regions, however, the filtering step refined detected associations by shifting peak locations and removing artefactual signals that would arise if genomic instability were not properly accounted for.

**Availability and implementation:** https://github.com/HautaniemiLab/SegmentQTL

## 1 INTRODUCTION

Quantitative trait loci (QTL) analysis identifies genetic variants associated with phenotypic traits (Aguet et al. 2023). In molecular QTL (molQTL) analysis, this approach is applied to phenotypes, such as gene expression levels (eQTL), by testing associations between genotypes and phenotypes within a defined genomic region surrounding the phenotype of interest.

When studying local regulatory effects, the analysis targets variants located near the gene or molecular feature, a process commonly referred to as cis-mapping. This approach tests associations between genotypes and phenotypes within a genomic window, typically spanning 100 kilobases (kb) to 1 megabase (Mb), to identify proximal variants that directly influence phenotype levels (Aguet et al. 2023). Variants significantly associated with phenotypes are classified as QTLs (Figure 1).

**Figure 1:**
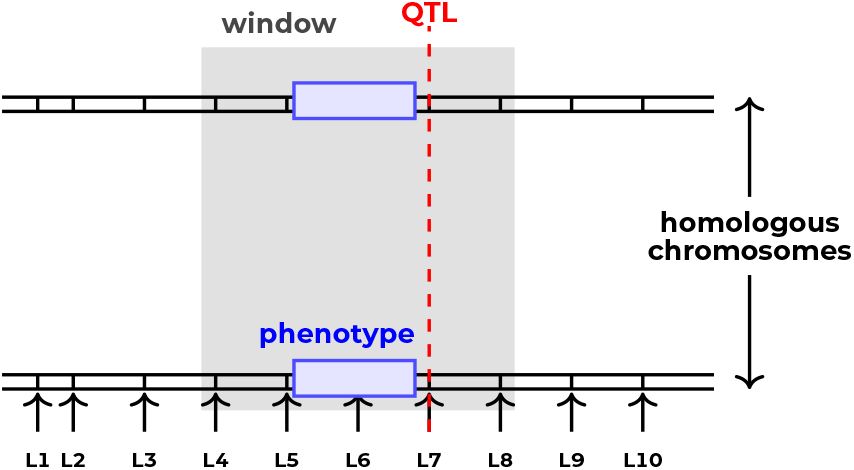
QTL mapping scheme, where the gray box illustrates the window in which associations are tested between the phenotype levels and allele dosages at variant loci, which are marked with arrows.

Copy number–driven cancers refer to specific type of cancers characterized by high genomic instability and extensive variation in gene copy numbers (Bhattacharya et al. 2020). Unlike in normal cells with a stable diploid status, in cancer cells the number of gene copies may vary from zero to several dozens. These alterations contribute to the development and progression of cancer. For example, missing copies can result in the loss of important tumor suppressor functions and amplifications of cellular oncogenes lead to uncontrolled activity of their downstream circuits (Jamalzadeh et al. 2024, Krais and Johnson 2020). Moreover, chromosomal breakpoints disrupt the spatial organization of the genome, dislocating regulatory elements, such as enhancers, from their target genes and thereby changing gene expression control.

Current molQTL analysis tools are not designed to operate in the complex genomic context of copy number–driven cancers. First, they typically assume discrete genotype classes (e.g., 0/1/2), which fail to capture the continuous dosage variation caused by aneuploidy and copy number alterations (CNAs) in cancer. Second, these tools test associations within fixed analysis windows without taking into account if structural changes have broken the link between variants and phenotypes. As a result, associations may be tested between loci that are no longer functionally connected, introducing false positives. To address both of these limitations, we developed SegmentQTL, a segmentation-aware molQTL method that supports continuous genotype dosages and filters out pairs of phenotypes and variants separated by breakpoints.

## 2 MATERIALS AND METHODS

### 2.1 SegmentQTL framework

SegmentQTL is a computational method designed to map molQTLs in cancers with frequent CNAs. It uses genomic segmentation data to improve the precision of molQTL mapping by accounting for chromosomal breakpoints, altered copy number, and continuous allele dosages.

#### 2.2.1 Variant loci selection

SegmentQTL uses segmentation information to filter variants within phenotype windows. If a phenotype and a variant within the theoretical analysis window are assigned to different segments, the genotype value is set as missing as illustrated in Figure 2. In addition, if a phenotype is located on more than one segment in a sample, all genotype dosages are set to missing in that sample for that phenotype-specific analysis window. This filtering is applied individually for each phenotype window to account for sample-specific segmentation patterns.

**Figure 2:**
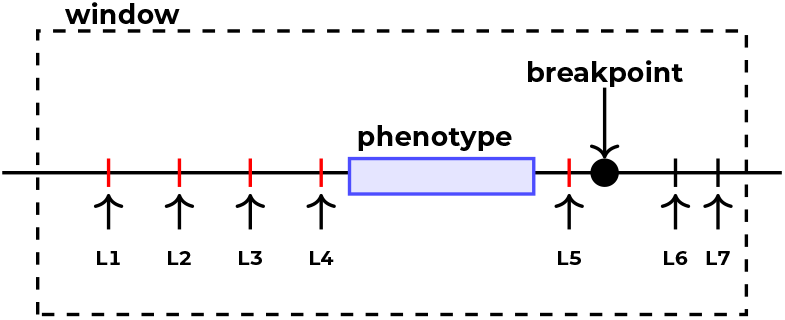
Variant loci selection in SegmentQTL mapping, where red variants (L1-L5) are included in the association testing whereas black variants (L6-L7) are filtered, because they are separated from the phenotype by a breakpoint.

#### 2.3.2 Association testing

SegmentQTL uses linear regression to find associations between genotypes and phenotypes, incorporating covariates to adjust for variant-independent effects. To isolate the effects of genotypes on phenotypes, we use residualization. The statistical approach is similar to the one used in FastQTL (Ongen et al. 2015) and tensorQTL (Taylor-Weiner et al. 2019) in cis-mapping.

Residualization is performed by fitting separate linear models for phenotypes and genotypes. Phenotypes and genotype dosages are each modeled as dependent variables in separate regressions, with covariates as independent variables. The residuals, representing variation unexplained by covariates, are then used as the inputs for downstream analyses.

We quantify the effect of genotypes by calculating the slope of the linear model and its standard error. The nominal *p*-value shows whether the slope of the relationship between residualized phenotype levels and residualized genotypes is significantly different from zero. It uses the squared t-statistic, Equation 1,

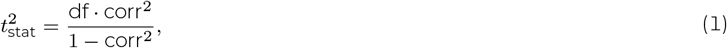

where *df* is the degrees of freedom and calculated as df = sample count − 2 − covariate count. 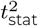 measures the strength of the correlation. It is used to calculate the *p*-value using the F-distribution as follows in Equation 2

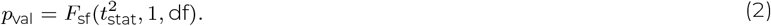

#### 2.3.3 Multiple testing correction

To improve computational efficiency and limit the number of required permutations to compute adjusted *p*-values, we apply a Beta approximation. This approach starts by estimating the *α* and *β* parameters of the Beta distribution, but instead of directly using the mean and variance of the permutation *p*-values, the parameters are refined iteratively. First, the degrees of freedom are optimized to ensure the shape parameter *α* of the Beta distribution closely matches the observed data. The optimization procedure matches the null distribution derived from permutation correlations to the Beta distribution.

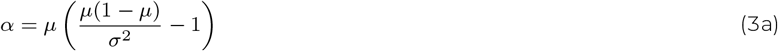

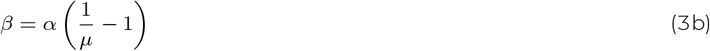

The initial estimates for *α* and *β*, Equation 3, are refined by minimizing the negative log-likelihood of the Beta distribution, as shown below in Equation 4:

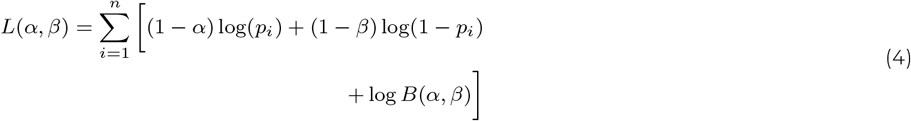

where *p*_*i*_ are the permutation-derived *p*-values, and *B*(*α, β*) is the Beta function. Once the parameters *α* and *β* are estimated, they are used to adjust the nominal *p*-values via the cumulative distribution function of the Beta distribution, Equation 5:

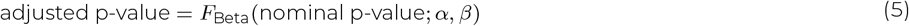

As an alternative to the Beta approximation, we implement a direct permutation scheme to adjust *p*-values. This approach permutes the whole dataset directly to empirically estimate the null distribution of correlations between phenotypes and genotypes. After generating the null distribution from permutations, the observed (nominal) correlation is compared against this distribution. The adjusted *p*-value is calculated as the proportion of permuted correlations with an absolute value greater than or equal to the absolute value of the observed correlation as seen in Equation 6:

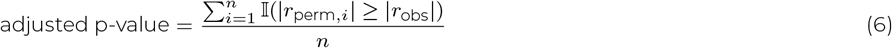

where *r*_perm,*i*_ represents the *i*-th permuted correlation, *r*_obs_ is the observed correlation, I is the indicator function, and *n* is the number of permutations.

False discovery rate (FRD) correction addresses the second levels on multiple testing correcting for the testing done for multiple phenotypes. For this purpose, SegmentQTL has a FDR mode, which uses Benjamini-Hochberg FDR-correction method.

#### 2.3.4 Covariate handling

Covariates were handled at two levels to control for confounding factors: sample-level and sample-phenotype-level. SegmentQTL treats copy number data as a sample-phenotype-level covariate and expects it as a required input, specifying copy number values for each phenotype in each sample.

In addition, SegmentQTL requires at least one sample-level covariate in numerical format. This ensures proper estimation of baseline expression levels across samples. If no biological covariates are available, a constant placeholder covariate (e.g. a vector of ones) can be used.

#### 2.3.5 Continuous genotypes

SegmentQTL supports continuous genotype dosage data, which is necessary for accurately capturing allele dosage variation in cancer. Unlike discrete genotype classifications, which assume a fixed number of allele copies, continuous dosages reflect the actual proportion of alleles present at a locus, accounting for copy number changes such as amplifications or deletions. In contrast, discrete genotypes would reduce this to a simple presence or absence call, losing critical quantitative information.

This continuous representation allows SegmentQTL to more faithfully reflect the underlying tumor biology, leading to more precise association testing and biologically meaningful results.

#### 2.3.6 Plotting functionality

SegmentQTL includes a built-in feature to generate molQTL plots for tested phenotype-genotype pairs, providing a visual representation of their relationships. These plots display genotype dosages on the x-axis and phenotype quantifications on the y-axis, grouped by dosage categories. This visualization allows observation of trends in the data and assessment of the statistical significance of associations, making the results more interpretable.

Users can control which phenotype-genotype pairs are plotted using the plot threshold parameter. This parameter allows users to set a threshold for selecting pairs to plot based on their statistical significance. For example, setting `plot_threshold` to one plots all tested pairs, while using a more stringent threshold (e.g., `plot_threshold` 0.05) restricts the output to only those pairs with *p*-values below the specified threshold.

#### 2.3.7 Parallelization

SegmentQTL can evaluate multiple phenotypes in parallel, utilizing multi-core processing to reduce runtime for large-scale analyses. It assigns one core per phenotype, enabling simultaneous analysis, and allows users to specify the number of cores. All cores specified must reside on the same machine or computational node.

### 2.2 DECIDER data

To demonstrate the functionality of SegmentQTL, we applied it to data from the DECIDER cohort. In the following, we outline the eQTL analysis workflow using DECIDER data, describing the preprocessing steps for SegmentQTL and the various data levels involved in the analysis (Figure 3).

**Figure 3:**
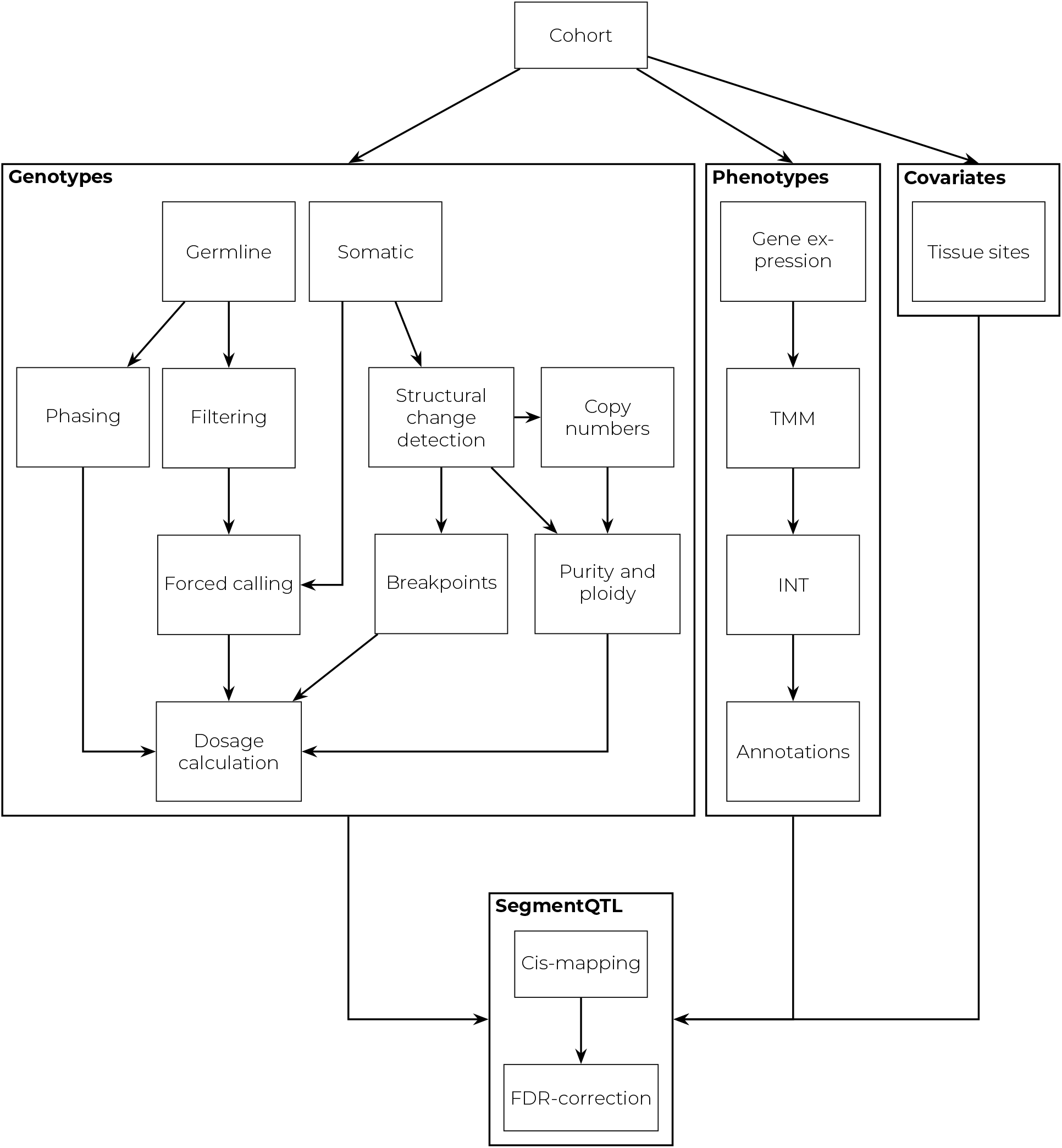
Workflow of the SegmentQTL analysis in DECIDER cohort, showing how genotype, phenotype, and covariate data were processed and integrated.

#### 2.2.1 Cohort description

The DECIDER cohort is a prospective, longitudinal study involving high-grade serous carcinoma (HGSC) patients treated with standard-of-care therapy at Turku University Hospital (ClinicalTrials.gov identifier: NCT04846933). As a copy number–driven cancer, HGSC exhibits frequent genomic rearrangements and structural alterations, creating a challenging genomic landscape for molQTL analysis. These characteristics makes the DECIDER dataset particularly suitable for testing SegmentQTLs ability to account for CNAs in molQTL mapping.

#### 2.2.2 Sample selection

DECIDER has collected patient samples from multiple cancer sites, including the primary site, metastases across peritoneal cavity and longitudinal samples from relapses. We included cancer samples with matching DNA and RNA sequencing data. For each patient, a single representative sample was selected to avoid bias from unequal sample numbers. We chose one treatment naïve sample with the highest CNA based purity (tumor fraction) for each patient. The final dataset consisted of samples from 199 patients.

#### 2.2.3 RNA sequencing phenotype data

We used gene expression levels derived from RNA sequencing data as phenotypes. The data preprocessing utilized the SePIA pipeline (Icay et al. 2016) within the Anduril framework (Cervera et al. 2019), as previously described in detail in Afenteva et al. 2024 and Häkkinen et al. 2021. Deconvolution using PRISM (Häkkinen et al. 2021) isolated the cancer cell component for downstream analysis.

Protein-coding genes with a median expression level exceeding 10 reads were retained for analysis. The data were normalized using edgeR (Chen et al. 2016), where trimmed mean of M-values (TMM) factors were calculated, and normalized counts were extracted using the cpm function. Inverse normal transformation (INT) was applied to transform the data into a Gaussian distribution using the RankNorm function from the RNOmni package (McCaw 2023). Finally, we obtained gene annotations, including gene start and end loci, from the Ensembl database (Martin et al. 2023).

#### 2.2.4 Whole-genome sequencing genotype data

Genotype data were derived from whole-genome sequencing (WGS) of cancer and peripheral blood samples. The germline genotypes of common biallelic polymorphisms (MAF *>* 0.05 in the cohort) were defined from the peripheral blood samples (Lahtinen et al. 2023). Hardy-Weinberg equilibrium calculations were performed using PLINK (Chang et al. 2015, Purcell and Chang n.d.), with a *p*-value threshold of 10^−8^ to exclude variants with significant deviations from expected allele frequencies.

Filtered germline variants were force called with GATK Mutect2 (Auwera and O’Connor 2020) in cancer sample sequencing data using the Anduril pipeline (Cervera et al. 2019).

Allele dosage was calculated by integrating variant read counts, tumor locus copy number, tumor purity, and phased genotypes. First, the allele depths were assessed at each variant site to determine the observed allele frequency. To account for copy number alterations, each variant was assigned segment-specific major and minor allele copy numbers, which, combined with tumor purity estimates, were used to compute the expected allele frequency. The alternate allele assignment was determined by comparing the observed and expected allele frequencies, selecting the configuration (major or minor allele) that best explained the observed data while considering the underlying CNV state. To improve assignment reliability, we incorporated haplotype phasing using SHAPEIT5 (Hofmeister et al. 2023) and the Sequencing Initiative Suomi (SISu) v4 reference panel from the Finnish Institute for Health and Welfare.

Once the alternate allele assignment was established, final dosages were calculated as the proportion of the assigned alleles copy number relative to the total copy number at the locus. In cases of uncertainty, local haplotype information and a smoothing procedure were applied to ensure consistency across neighboring variants. The allele dosage calculation script is available at https://github.com/HautaniemiLab/AlleleDoser.

#### 2.2.5 Segmentation, copy number, and purity data

Copy number segmentation was analyzed using a combination of GRIDSS and the Hartwig Medical Foundation (HMF) toolkit (https://github.com/hartwigmedical/hmftools, Micoli et al. manuscript in preparation). Breakpoints were identified with GRIDSS (v2.13.2) (Cameron et al. 2021), excluding regions listed in both the ENCODE (Amemiya et al. 2019) and in-house DECIDER blacklists (Lahtinen et al. 2023). Somatic filtering was performed with GRIPSS (v2.0) (Cameron et al. 2019), utilizing a panel of normals derived from blood samples of DECIDER patients and a Dutch population. B-Allele Frequency (BAF) was calculated using the AMBER tool (v3.8) with heterozygous biallelic loci identified by GATK (Auwera and O’Connor 2020). Read depth with GC-content normalization was performed using COBALT (v1.12). The combined outputs from these tools, along with somatic single nucleotide variants identified by GATK as described in Lahtinen et al. 2023, were used in PURPLE (v3.7.2) (Cameron et al. 2019, Priestley et al. 2019) for the estimation of copy number segmentation profiles, tumor purity, and ploidy for the analyzed samples.

#### 2.2.6 Covariates

Sample-level covariate included the tumor site to account for location-specific gene expression differences. Gene-specific copy numbers, corrected for ploidy by dividing raw copy number values by the ploidy estimate, were used as sample-phenotype-level covariates. The correction ensured that the covariate reflects gene dosage rather than absolute copy number.

#### 2.2.7 Gene set enrichment analysis

We performed gene set enrichment analysis (GSEA) using the fgsea R-package (Korotkevich et al. 2019) to identify enriched pathways among the association results. Genes were ranked by their slope values from the linear models used in the molQTL analysis, and gene sets from the Hallmark collection of the Molecular Signatures Database were retrieved via the msigdbr R-package (Dolgalev 2022).

#### 2.2.8 GISTIC analysis

We ran GISTIC 2.0.23 (Mermel et al. 2011) with default parameters to identify regions of recurrent copy number alterations in the segmented copy-number data (see Section 2.2.5).

## 3 RESULTS AND DISCUSSION

### SegmentQTL: molQTL analysis in copy number–driven cancers

SegmentQTL is a computational method that identifies variants associated with molecular phenotypes in copy number–driven cancers. In contrast to current molQTL analysis tools, SegmentQTL incorporates genomic segmentation data to enhance precision and biological significance. By considering segment boundaries, we avoid artifacts arising from testing of variant-gene pairs which have lost their physical contact in a somatic structural aberration.

### Applying SegmentQTL to HGSC

SegmentQTL is particularly well suited to cancers with extensive genomic instability, such as HGSC. When applied to HGSC tumor data, SegmentQTL identified 6370 eGenes whose expression was significantly associated with nearby tumor germline variants. We define a gene as an eGene if its FDR-corrected, permutation-adjusted *p*-value falls below 0.005.

The nominal analysis revealed genomic regions with significant variant-gene associations across the genome. The Manhattan plot alongside a GISTIC track highlighting recurrent CNAs in Figure 4 visualizes the distribution and strength of these associations. We identified strong eQTL associations in both genomically stable regions with few CNAs and in unstable regions characterized by frequent alterations. These findings show that significant associations can be detected across the genome, but they also raise the question of how often genomic instability interrupts variant-gene associations.

**Figure 4:**
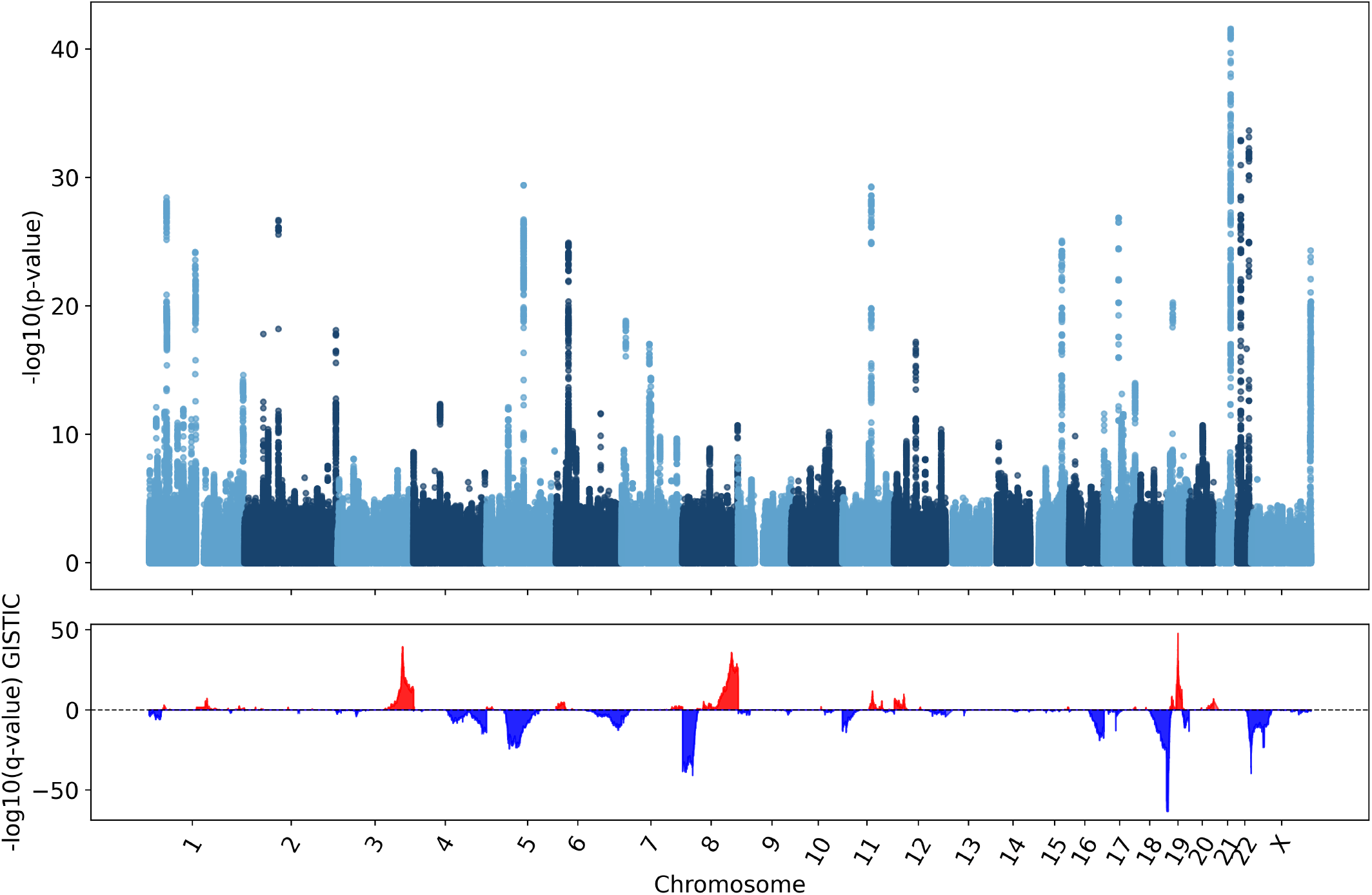
Manhattan plot of nominal analysis results with GISTIC track highlighting regions of recurrent copy number alterations. Significant associations are detected both from extensively altered genomic regions and more quiet regions.

We assessed how breakpoints affect molQTL analysis by examining changes in effective sample size across the analysis window for each phenotype. The effective sample size represents the number of samples in which the variant and the phenotype remain within the same continuous genomic segment. As shown in Figure 5, large changes in effective sample size are common: even in genomically stable regions, at least 10% of samples typically contain a breakpoint within the analysis window. In contrast, even in unstable regions with frequent chromosomal breaks, more than half of the samples often retain intact variantphenotype pairs.

**Figure 5:**
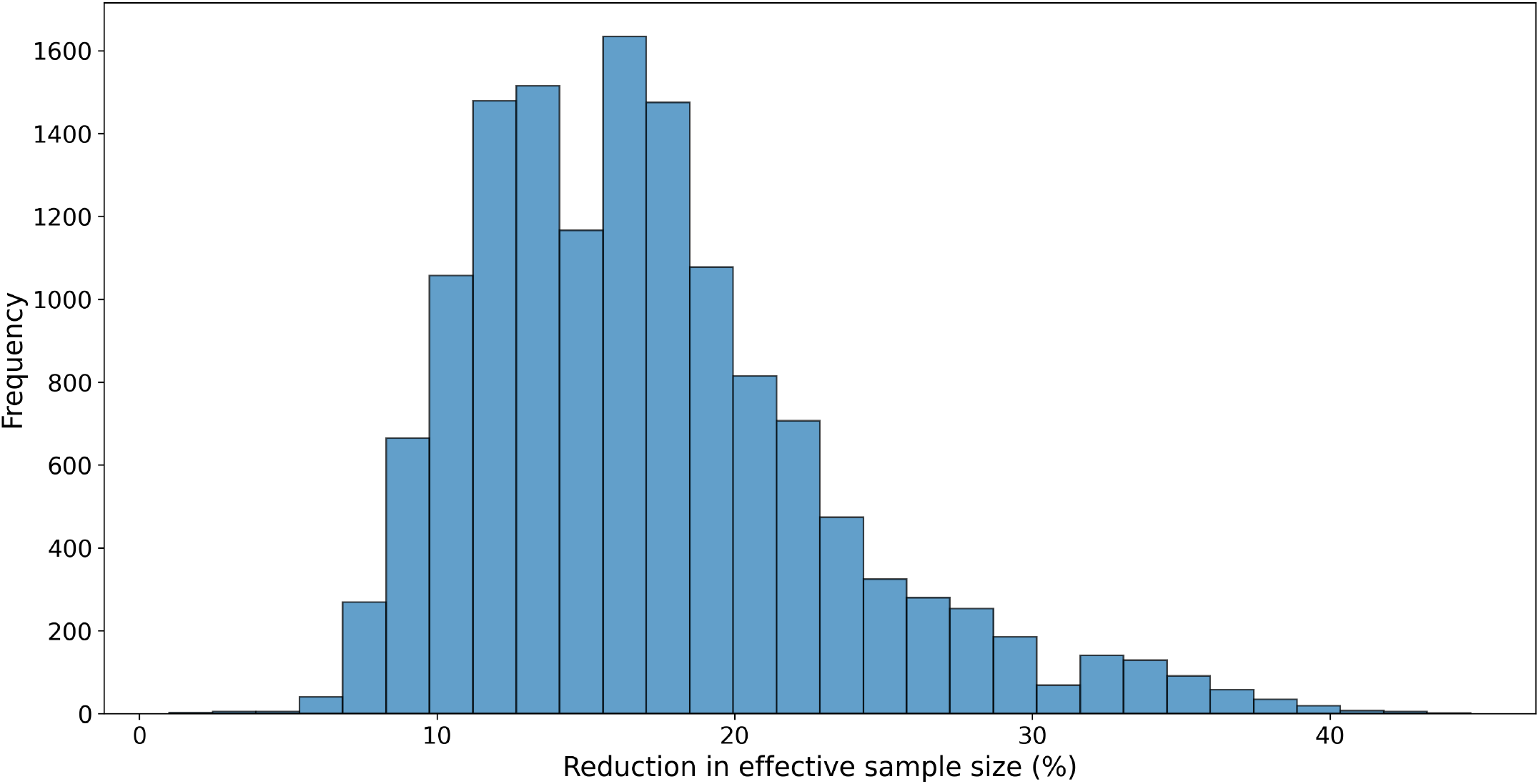
Distribution of maximum differences in effective sample sizes within molQTL analysis windows, demonstrating the impact of filtering by genomic segments.

These findings emphasize that breakpoints are widespread and that their positions vary considerably across patients. Incorporating segmentation-aware filtering is therefore important to avoid testing disrupted associations.

To better understand the affected oncogenic processes, we performed GSEA. Among the five significantly enriched Hallmark pathways (Table 1) were the epithelial-mesenchymal transition and regulation of the KRAS signaling pathway. While these pathways have frequently been implicated in the HGSC pathogenesis, via acquisition of somatic aberrations (Handley et al. 2022, Jamalzadeh et al. 2024), our results suggest the selection of germline regulatory alleles is an important oncogenic mechanism.

**Table 1:** Top enriched Hallmark pathways (adjusted *p*-value *<* 0.01) from GSEA results with corresponding normalized enrichment scores (NES).

| Pathway | padj | NES |
| --- | --- | --- |
| Epithelial mesenchymal transition | $3.26 \times 10^{-3}$ | -1.88 |
| Estrogen response late | $3.26 \times 10^{-3}$ | 1.84 |
| Complement | $3.39 \times 10^{-3}$ | 1.85 |
| KRAS signaling (down) | $6.05 \times 10^{-3}$ | 1.95 |
| Apical surface | $6.11 \times 10^{-3}$ | 2.07 |

Finally, we evaluated the computational performance of SegmentQTL in two analysis modes: nominal analysis with all variants and permutation analysis using the beta approximation method with 8000 permutations. All analyses were conducted on a 12th Gen Intel^®^ Core^™^ i5-1235U processor. In the nominal analysis mode, the median runtime per phenotype was 11.59 seconds (SD. 4.30 seconds). For the permutation mode, the median runtime per phenotype was 29.52 seconds (SD. 2.44 seconds). These runtimes reflect only the actual processing time and do not include input file reading or the output writing.

### Evaluating SegmentQTL’s performance against existing method

To assess the performance of SegmentQTL, we compared the results with tensorQTL (Taylor-Weiner et al. 2019), a previously published state-of-the-art method for molQTL analysis. The evaluation focused on selected genes from both a genomically stable and an unstable region.

The first part of our evaluation focused on nominal analysis with all variants. In a stable chromosomal region with only few break-points, Figure 6A, SegmentQTL produced results highly similar to tensorQTL, with matching peaks observed for genotype-phenotype associations. Here, we define peaks as the genomic loci with the smallest *p*-values, separated by a minimum distance of 50,000 base pairs. The similarity demonstrated that in the absence of genomic instability, both methods effectively capture eQTL signals. However, in an unstable region with frequent chromosomal breaks, Figure 6B, the results diverged due to SegmentQTLs integrated purifying filtering step. This step removed associations where the gene and variant are separated by a breakpoint in a given sample. As a result, the detected peak locations and associated variants differed between the tools. These differences were not just technical, but they may lead to focusing on variants that are not truly regulatory or to incorrect classification of a gene as an eGene or not.

**Figure 6:**
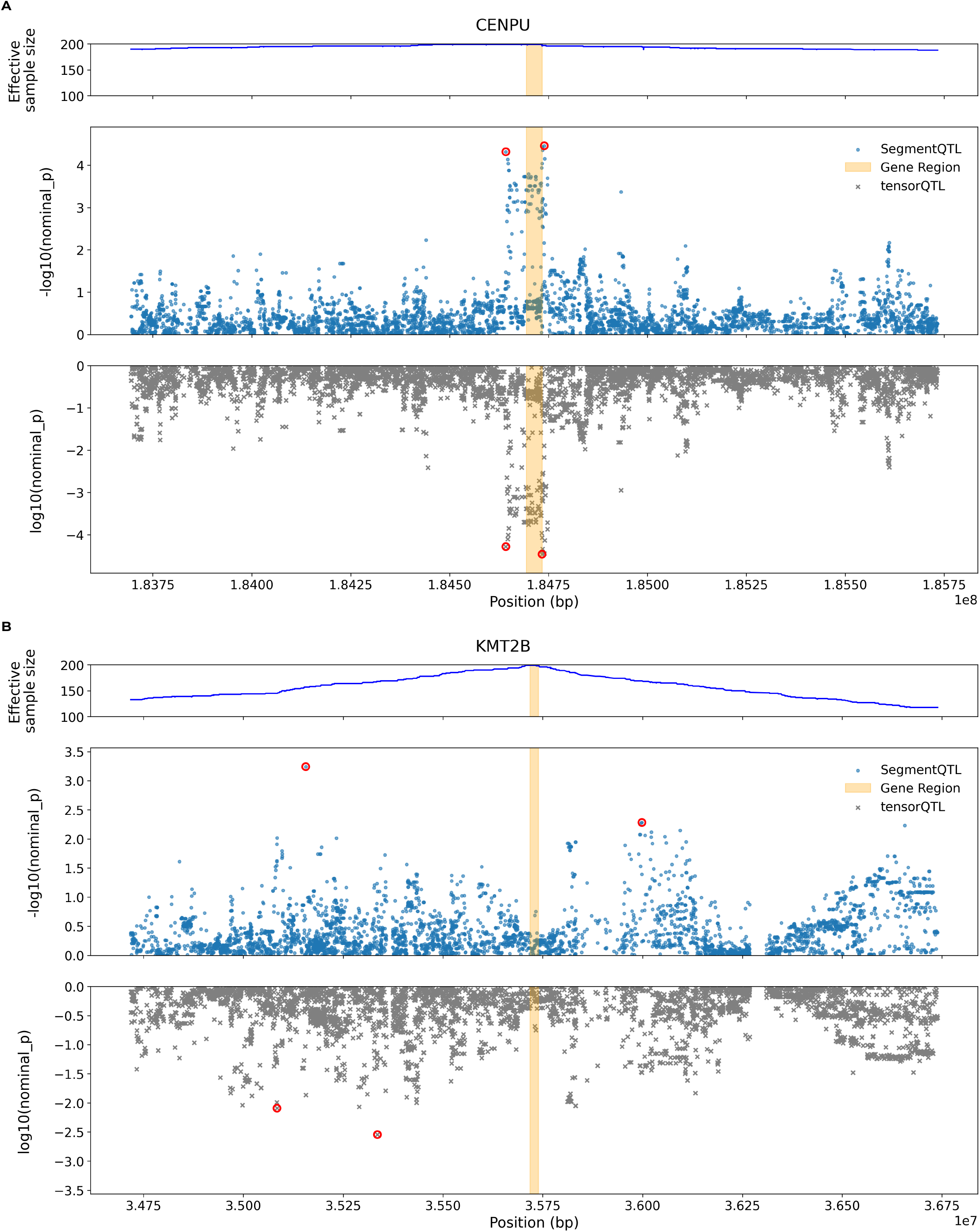
Comparison of SegmentQTL and tensorQTL results for selected genes in stable (A) and unstable (B) genomic regions, highlighting the peak locations in red. In the stable region, both methods identify similar background signals and peaks. In contrast, results diverge in the unstable region due to SegmentQTLs by-segment filtering, as reflected also in the effective sample size track.

Across chromosomes, the median concordance of nominal *p*-values between SegmentQTL and tensorQTL, measured by Spearman’s rho, was 0.85 (s.d. 0.03, IQR 0.04). The concordance was consistent across most chromosomes except for chromosome 19, which was an outlier with the lowest concordance of 0.74. This discrepancy was likely due to the high level of aneuploidy in chromosome 19, which may introduce variability in the eQTL signals between the two methods.

Upon conducting permutation testing, we identified major differences between SegmentQTL and tensorQTL. tensorQTL failed to execute permutation analysis with continuous-valued genotypes, halting the process unless the genotypes were rounded to integers. This rounding step, however, mitigated the aneuploid dosage variation characteristic to cancer. As a result, tensorQTL identified only 63 eGenes, which was not only substantially lower than the number detected by SegmentQTL but also considerably below what is generally expected for eQTL analyses with this sample size (Aguet et al. 2023) when using a similar FDR correction method and threshold, demonstrating the need for a tailored methodology for analysis of copy number-driven cancers.

While SegmentQTL is specifically designed for cancer-related datasets, its advantages are context-dependent. In datasets with limited copy number variation, such as normal tissues or genomically stable regions, SegmentQTL does not offer clear advantages over existing tools. In such cases, the additional segmentation-aware filtering step may only increase computational runtime without improving performance. Nevertheless, in the various types of copy number–driven cancers, including triple-negative breast cancer, lung cancer, or esophageal cancer (Steele et al. 2022), currently lacking valid tools for molQTL analysis, SegmentQTL opens completely unexplored avenues for improved understanding of the consequences of allelic selection along cancer progression.

## 4 CONCLUSION

Molecular QTL analysis is a powerful tool for studying how genetic variants influence molecular phenotypes. However, existing methods do not account for the high genomic instability characteristic of copy number–driven cancers, making them unsuitable for such studies. To address this limitation, we developed SegmentQTL, a molQTL analysis method that incorporates sample-specific segmentation information.

SegmentQTL applies an integrated purifying filtering step to eliminate misleading signals produced by extensive structural changes in the genome. Without this filtering, associations disrupted by chromosomal breaks would still be included in the analysis, increasing background noise and leading to false discoveries. By excluding variants that are separated from the tested phenotype by a breakpoint, SegmentQTL enhances the accuracy and biological relevance of the cis-mapping results. This improved resolution may aid in the identification of candidate biomarkers and therapeutic targets in cancers with high genomic instability.

## ACKNOWLEDGMENTS

We are deeply grateful for all the individuals who took part in the DECIDER study and all the clinicians, technicians and administrative staff who have enabled this work to be carried out, especially the DECIDER lead clinician Dr Johanna Hynninen, and the research coordinators Dr Ann-Christin Ostwaldt and Elina Valkonen.

The data used for the research was phased with the THL Biobanks SISu v3 reference panel obtained from THL Biobank. We thank all study participants for their generous participation in the FINRISK, Health 2000 and Migraine Family studies. We also thank the Sequencing Informatics Team, FIMM Human Genomics, University of Helsinki for the work done in preparation of the reference panel data.

Computing resources were provided by CSCIT Center for Science Ltd..

D.A. gratefully acknowledges support from the Orion Research Foundation.

## DATA AVAILABILITY

Whole genome sequence data and transcriptome sequencing data has been deposited at the European Genome-phenome Archive (EGA), which is hosted by the EBI and the CRG, under the respective accession numbers EGAS00001006775 and EGAS00001004714.

